# Comparison of software packages for detecting unannotated translated small open reading frames by Ribo-seq

**DOI:** 10.1101/2023.12.30.573709

**Authors:** Gregory Tong, Nasun Hah, Thomas F. Martinez

## Abstract

Accurate and comprehensive annotation of microprotein-coding small open reading frames (smORFs) is critical to our understanding of normal physiology and disease. Empirical identification of translated smORFs is carried out primarily using ribosome profiling (Ribo-seq). While effective, published Ribo-seq datasets can vary drastically in quality and different analysis tools are frequently employed. Here, we examine the impact of these factors on identifying translated smORFs. We compared five commonly used software tools that assess ORF translation from Ribo-seq (RibORFv0.1, RibORFv1.0, RiboCode, ORFquant, and Ribo-TISH), and found surprisingly low agreement across all tools. Only ∼2% of smORFs were called translated by all five tools and ∼15% by three or more tools when assessing the same high-resolution Ribo-seq dataset. For larger annotated genes, the same analysis showed ∼72% agreement across all five tools. We also found that some tools are strongly biased against low-resolution Ribo-seq data, while others are more tolerant. Analyzing Ribo-seq coverage as a proxy for translation levels revealed that highly translated smORFs are more likely to be detected by more than one tool. Together these results support employing multiple tools to identify the most confident microprotein-coding smORFs, and choosing the tools based on the quality of the dataset and planned downstream characterization experiments of predicted smORFs.

## INTRODUCTION

Early efforts to annotate eukaryotic genomes relied in part on applying expected properties of coding regions, such as having an ATG/AUG start codon in frame with a downstream stop codon, one protein coding region per transcript that is often the longest open reading frame (ORF), and a minimum length cutoff of 100 codons to identify overlooked coding regions [1]. While effective, there remained the possibility that ORFs which do not follow these rules can be translated to encode functional proteins. Recent advances in genomics, proteomics, and bioinformatics have allowed researchers to empirically define protein coding regions within genomes with better precision. The most striking result of these new studies is that thousands of small open reading frames (smORFs) containing less than 100-150 codons, which were presumed to be randomly occurring and non-functional, are in fact translated into small proteins dubbed microproteins. These smORFs make up the majority of unannotated ORFs and represent an increasingly active area of research. Many microproteins have now been shown to be critical in normal biological processes and disease.

One of the primary methods for re-annotation of genomes is based on ribosome profiling (Ribo-seq). Ribo-seq involves stalling elongating ribosomes in cell or tissue lysates with the small molecule inhibitor cycloheximide, followed by digestion of polysomes with an RNase and preparation of the ribosome protected RNA fragments (RPFs) into next generation sequencing libraries. Following sequencing, the resulting reads are processed and aligned to the genome to determine the locations of the ribosomes in each sample at harvesting. By identifying the locations of ribosomes, bioinformatic tools can then be applied to infer which open reading frames are translated. However, due to the variation in Ribo-seq protocols and a variety of different software tools that have been developed to analyze translation from Ribo-seq data, there is no consensus on best practices within the field for predicting smORFs.

For the field to progress further toward functional investigation of individual microproteins and exploration of their utility as therapeutic targets, confidence in which smORFs are annotated as translated is needed. Previously, we showed that differences in Ribo-seq data quality can strongly impact which smORFs are called translated and that analyzing biological replicate datasets is helpful for separating robustly translated smORFs from noise [2]. Here, we hypothesized that different software tools for interpreting Ribo-seq data can also introduce inconsistencies into which smORFs are considered translated due to differences in the properties of Ribo-seq data are considered in scoring, how they are weighted, and what statistical methods or classifiers are applied. To understand how the choice of software tool can influence smORF prediction, we evaluated the performances of several popular Ribo-seq-based ORF prediction tools. We found that while all tools show high congruence when identifying larger annotated ORFs as translated, they show low similarity for which unannotated smORFs are predicted to be translated. Analysis of Ribo-seq coverage levels between annotated ORFs and unannotated smORFs suggest that the overall lower translation levels of smORFs contributes to their noisier translation predictions. In addition, we observed large differences between the tools’ abilities to predict smORF translation when using lower quality Ribo-seq datasets versus high. We also demonstrated that incorporation of an RNA-seq-derived *de novo* transcriptome assembly can add additional unannotated smORFs compared to using a standard GENCODE transcriptome annotation. Altogether, these results highlight the importance of using multiple tools to raise confidence in the annotation of individual ORFs for functional studies and broaden the pool of potential smORFs to test in high-throughput screens.

## METHODS

### Ribo-seq datasets and preprocessing

Ribo-seq datasets analyzed in this study were generated in our previous study [3], and can be downloaded from the Gene Expression Omnibus (GEO) database repository under accession number GSE125218. The specific Sequence Read Archive (SRA) IDs for the Ribo-seq datasets are as follows: high-resolution HeLaS3 - SRR8449578, low-resolution HeLaS3 -SRR8449575, harringtonin (TI-seq) HeLaS3 - SRR8449585, high-resolution HEK293T - SRR8449568, medium-resolution HEK293T - SRR8449567, and low-resolution HEK293T - SRR8449566.

Ribo-seq reads were preprocessed by trimming of 3’ adapter sequences (AGATCGGAAGAGCACACGTCT) using the FASTX-toolkit. Next, reads aligning to rRNA and tRNA sequences were filtered out using STAR with parameters –outReadsUnmapped Fastx and the remaining reads were subsequently aligned to the GENCODE hg38 version 39 genome assembly using STAR with the following settings –outFilterMismatchNmax 2 – outFilterMultimapNmax 4 –chimScoreSeparation 10 –chimScoreMin 20 –chimSegmentMin 15 – outSAMattributes All –outSAMtype BAM SortedByCoordinate. The resulting bam file was filtered for primary alignments using samtools with the following parameters -bS -F 0X100. After, multimappers were removed using samtools with the following parameters -bq 255. The alignment files used for RiboCode’s prepare_transcripts function requires the use of the quantMode option during STAR alignment. To run RiboCode, reads were processed separately using author recommended settings to include —outfilterMismatchNmax 2 –outSAMtype BAMSortedByCoordinate –quantMode TranscriptomeSAM Genecounts – outFilterMultiMapNmax 1 –outFilterMatchNmin 16 –alignEndsType EndToEnd. Length histograms were generated by sampling a million reads and sorted by length from the final alignment file. Metagene plots were created using RibORFv0.1’s readDist.pl function and a custom script was used to calculate the fraction of in frame reads based on the total corrected reads. Other tools also have the capability to generate metagene plots. To ensure the same set of read lengths were used for analysis across the different workflows, the same read lengths and offset corrections were used for all ORF predictions for each separate library. Ribo-seq coverage was visualized by generating bedgraphs using HOMER and uploading the bedgraphs to the UCSC Genome Browser.

For final list filtering, smORFs with a minimum length cutoff of >6 amino acids and maximum length cutoff of 150 amino acids was applied to all smORF lists. Afterwards, bedtools intersect was used to remove smORFs that had over 90% overlap with CDS regions of canonical genes with the following commands, -f 0.9 -v -s. We chose to exclude smORFs that overlap fully with annotated ORFs in our analysis as they can be difficult to accurately identify by Ribo-seq, but all the tools will allow for fully internal smORFs to be scored. After filtering out passing smORFs, an additional filter using BLASTP was applied to remove potential pseudogenes and potentially missed RefSeq annotated microproteins. The settings for running the BLASTP search was -outfmt 10 -max_target_Seqs 5 -evalue 0.0001, and microproteins with BLASTP scores ≥40 were filtered out. For generating translation scores for annotated genes, RibORFv0.1 was run using a separate refFlat containing GENCODE CDS regions. For RiboCode, Ribo-TISH, ORFquant, and RibORFv1.0, annotated genes that were detected were separated out from the final list of ORFs predicted.

### Tools compared in this study for microprotein-coding smORF identification

#### RibORFv0.1

RibORFv0.1 is the oldest tool of those we compared and is the tool we have used to annotate microprotein-coding smORFs in our previous studies [2,4]. RibORFv0.1 utilizes a support vector machine classifier to select for translating ORFs based on fraction of A-site reads aligned to the correct reading frame and read distribution uniformity over the ORF. The model uses canonical ORFs and off-frame ORFs for positive and negative controls, respectively, to train the classifier to predict smORFs. A final p-value score is determined based on these two properties. The authors suggest a score of ≥0.7 as a threshold for translation. Importantly, this tool requires the user to provide a list of ORFs to be scored and cannot use the Ribo-seq data to help identify start and stop sites. ORFs were defined using a custom java script, GTFtoFASTA [2]. Using the reference GENCODE transcriptome, all three open reading frames were parsed to find the most upstream canonical ATG start codon and in frame stop. If there is no canonical start codon, then the ORF is defined from stop codon to stop codon. Running RibORFv0.1 for translation scoring, ORFs were filtered with a minimum length cutoff of 18 and minimum read coverage cutoff of 10 using the ribORF.pl script. The resulting list of ORFs was further filtered with a pvalue cutoff of ≥0.7, max nucleotide length cutoff of 450, and a read coverage cutoff of 10.

#### RibORFv1.0

RibORFv1.0 is an updated version of RibORF that uses a different strategy for scoring translation but is otherwise similar to RibORFv0.1. Instead of a support vector machine classifier, RibORFv1.0 uses a logistic regression model to determine the pvalue scores. In addition, RibORFv1.0 no longer uses a pre-scored training set of known translated and non-translated ORFs but uses the user’s own data to train prediction parameters based on pre- defined positive and negative ORFs. It also parses user provided transcriptomes to identify all possible ORFs and thus does not require a user provided list. ORF scoring was processed by first running the ORFannotate.pl script with default settings. After candidate ORFs are generated, ribORF.pl was used to identify translated ORFs using default settings of orfLengthcutoff of 6 and readlengthcutoff 11. As with RibORFv0.1, the scored ORF list was filtered with a pvalue cutoff of ≥0.7 and max nucleotide length cutoff of 450.

#### Ribo-TISH

Like other tools, Ribo-TISH can assess ORFs for translation using standard Ribo-seq data from samples treated with cycloheximide. In addition, it can use translation initiation sequencing (TI-seq) data from cells treated with translation initiation inhibitors, e.g. harringtonin or lactimidomycin, to identify translated ORFs either with TI-seq data alone or in combination with Ribo-seq data. For scoring, it uses a non-parametric Wilcoxon rank-sum test for its assessment of 3-nt periodicity. Ribo-TISH can also parse user provided transcriptomes to identify all possible ORFs and *de novo* annotate the translatome. Ribo-TISH was run with the strategy of taking the most distal start codon to stop codon with RPF coverage when defining the ORF. The predict function with the parameters –longest –altcodons TTG,CTG,GTG –seq – aaseq with a p-value threshold of <0.05 was used. For Ribo-TISH analysis with translation initiation data, the same settings were used with the additional -t flag for the harringtonin dataset input.

#### RiboCode

RiboCode is a *de novo* translatome annotation software that relies solely on the 3-nt periodicity pattern. For scoring, RiboCode uses a modified Wilcoxon signed rank-sum test to assess whether the P-site density for a particular ORF is greater than the densities in the alternative reading frames. Like the other modern tools, RiboCode parses a user provided transcriptome to identify all possible ORFs for scoring. RiboCode also allows for the user to input non-canonical start codons to use for defining candidate ORFs. Detection of translated ORFs was identified using the RiboCode function with the settings -l no -s ATG -A CTG,GTG,TTG -g and the default p-value cutoff of 0.05.

#### ORFquant

ORFquant is also able to *de novo* annotate the translatome. It uses a multitaper test to select in-frame signal showing 3-nt periodicity, similar to the older RiboTaper tool developed by the same author. This tool generates a p-value and a cutoff of 0.05 is used to identify translated ORFs. Importantly, ORFquant requires the average signal on each covered codon to be >50% in frame and only considers AUG start codons. ORFquant was run using the authors’ recommended settings. First, a .2bit file and gtf are used to create a TxDb and Rdata file using the prepare_annotation_files function. Next, the prepare_for_ORFquant function was used to process the alignment bam file and text file containing the read lengths and cutoff for analysis. Lastly, run_ORFquant was used to take the files produced in the previous steps to score ORFs using a p-value threshold of 0.05.

### Ribo-seq Read Coverage and PhyloCSF Analysis

Ribo-seq read coverage for candidate smORFs identified by each tool were quantified alongside the top expressed isoforms for annotated genes. Coverage was quantified using HOMER’s analyzeRepeats function and expression was normalized by transcripts per million (TPM). Average PhyloCSF scores for the 58-mammal alignment used with genome build hg38 were extracted for all smORFs from the UCSC genome browser’s PhyloCSF Track Hub.

### Nanopore Long-read library preparation and sequencing

Total RNA was isolated from HeLa-S3 using the QIAGEN RNeasy kit. RNA integrity was assessed using TapeStation 4200 (Agilent) and RNA samples with RIN> 8 were used for library preparation for long read sequencing. Isolated total RNA was used to generate sequencing library following Oxford Nanopore Technologies protocol for cDNA-PCR sequencing kit. 50 ng of total RNA was first reverse transcribed for complementary strand synthesis using strand switching primers. cDNA was PCR amplified using primers that contain 5’ tags, which enables attachment of rapid sequencing adapters. The cDNA library was loaded onto R9.4.1 flow cells according to Oxford Nanopore Technologies protocol and sequenced for 48 hours with High accuracy setting on GridION system in the Salk NGS core.

### De novo transciptome assembly

For the long read RNA-seq datasets generated using the Nanopore sequencing platform, reads were processed using the FLAIR pipeline. Reads were aligned using FLAIR align module with minimap2 and converted to a SAM file in BED12 format. FLAIR correct was used to correct misaligned splice sites using the GENCODE version 39 annotation. Finally, FLAIR collapse takes the high confidence isoforms from the corrected reads to output a gtf. Using the StringTie merge option, the FLAIR gtf was merged with the GENCODE reference gtf to create a combined non-redundant set of transcripts used for downstream analysis.

For the paired-end RNA-seq datasets generated using the Illumina sequencing platform, originally generated in [2], fastq files were downloaded from the SRA with accession codes found in (Table S1) and trimmed of adapter sequences using TrimGalore. Reads were aligned using STAR with the options –runMode alignReads –sjdbOverhang 100 –runRNGseed 133 – twopassMode Basic –outSAMstrandField intronMotif –outfilterINtronMotifs Remove Noncanonical –outSAMattributes All. The resulting bam file was then sorted using samtools. For each library, StringTie was used to assemble transcripts from the sorted bam files using the guided assembly option. The assembled transcripts were then merged using the StringTie merge option with the GENCODE reference transcriptome annotation. The resulting gtf file was used as the transcriptome for downstream smORF analysis. GFFCompare was used to compare and evaluate the two transcriptome assemblies.

## RESULTS AND DISCUSSION

### Tools for detecting translated open reading frames from Ribo-seq

We compared five popular tools for analyzing individual open reading frames for translation using Ribo-seq data, including RibORF version 0.1 (RibORFv0.1) [5], RibORF version 1.0 (RibORFv1.0) [6], Ribo-TISH [7], RiboCode [8], and ORFquant [9]. These tools were published between 2015 and 2020 and have been applied frequently to identify novel translated ORFs, including smORFs, in the years since. Each tool includes an assessment of the 3 nucleotide (3-nt) periodicity of aligned ribosomal A-site or P-site reads that are in-frame with a particular ORF to aid in scoring translation. This feature is a hallmark of active translation as the ribosome scans ORFs translating 3-nt codons from the start codon to the stop codon [10].

Higher resolution datasets have a higher percentage of reads in-frame with annotated ORFs. However, the statistical methods applied for assessing whether the fraction of in-frame reads is significant differs widely. For example, RibORFv0.1 utilizes a support vector machine approach to classify and score ORFs, while RiboCode uses a modified Wilcoxon signed-rank sum test to determine the significance in-frame versus out-of-frame read enrichment within the tested ORF **(Fig. 1A)**. In addition, whether tools allow for ORFs initiating from near-cognate start codons, such as CUG or GUG, or consider other features such as percent ORF coverage differs among the tools. More details on how each tool scores translation is included in the Methods section.

**Figure 1.**
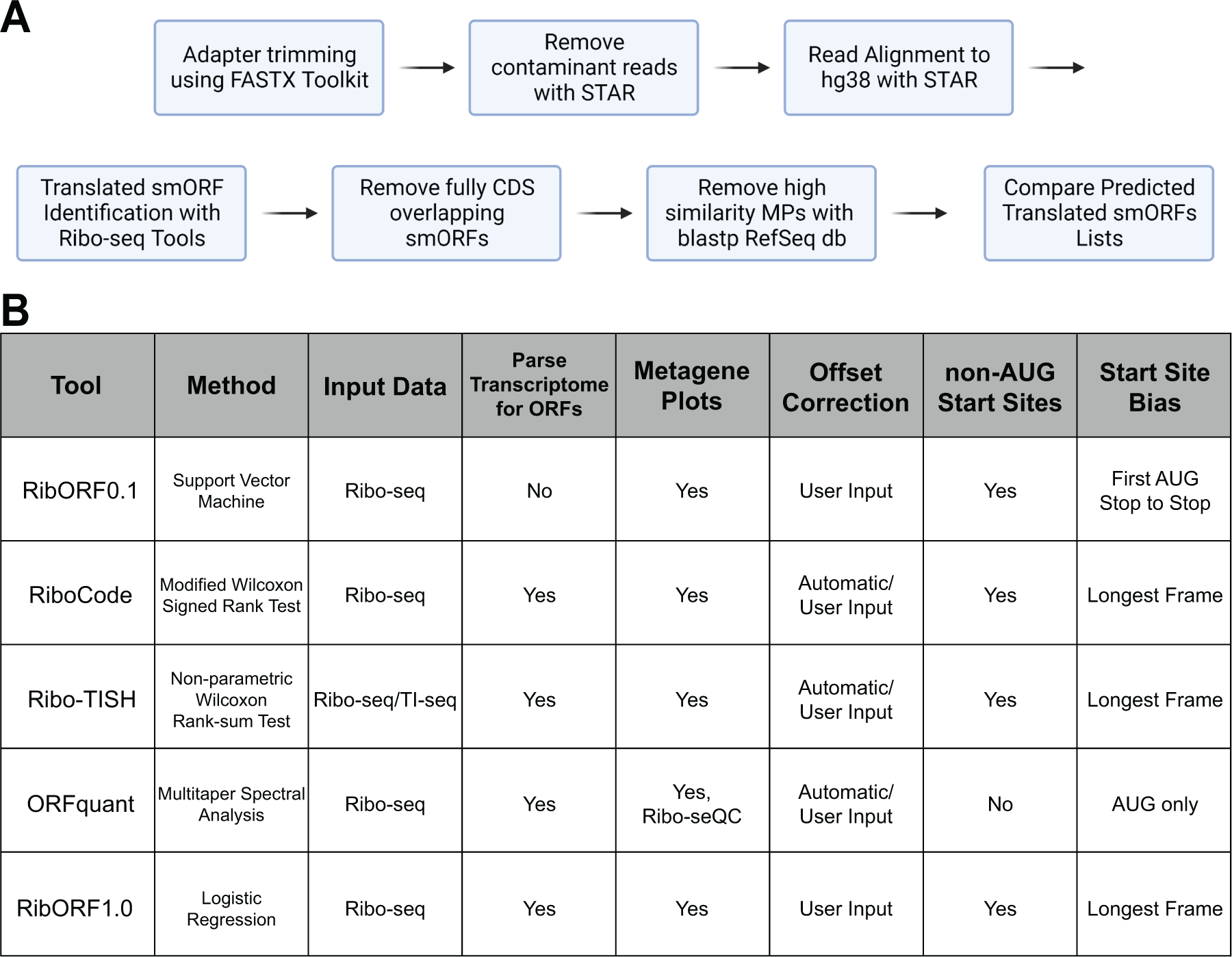
Workflow of smORF Annotation and Ribo-seq Tool Features. (A) Workflow for processing and filtering of Ribo-seq datasets that were used for ORF identification and comparison of translated unannotated smORF lists. **(B)** Properties of the computational methods compared in this study.

To compare the tools, we developed a standardized workflow to take unprocessed Ribo-seq data and generated a filtered list of predicted novel smORFs **(Fig. 1B)**. To summarize, 3’- adapter sequences are trimmed and reads aligning to rRNA and tRNA sequences are filtered out. The remaining reads are mapped to the hg38 genome using STAR [11] and the resulting alignment file of only uniquely mapped reads is used as the input for each tool to score ORFs for translation. Each tool was also given either a list of all possible ORFs to score, which we generated from the GENCODE comprehensive set of human transcripts, or the entire GENCODE transcriptome file for the software to parse into ORFs for scoring. The Ribo-seq datasets analyzed in our tool comparison were generated in our previous study [2] and include low- and high-resolution datasets collected from HeLa-S3 and HEK293T cell lines (**Supplementary** Fig. 1). The high-resolution datasets show greater than 70% in-frame RPF read alignment with known coding regions across all read lengths retained for analysis, while low-resolution data show only ∼50% of in-frame RPF reads (**Supplementary** Fig. 2). These datasets allowed us to assess any differences between the tools in handling varying quality data and ensure that any observed trends are not cell line specific. Following scoring by each tool, smORFs that were found to fully overlap within annotated CDS regions were removed. These internal smORFs can be difficult to accurately score by Ribo-seq as reads aligned to each ORF inherently lowers the score of the other ORF. The list of remaining unannotated smORFs were then used for comparison across tools.

### Comparing Predicted Translated smORFs Across Tools

In our previous study, we showed that there was a high overlap in the detection of annotated coding regions from Ribo-seq data across different resolutions, but that the list of smORFs called translated was noisy and showed low overlap across datasets [2]. This study only used RibORFv0.1 to analyze smORF translation, leaving an open question as to whether the poor overlap was an artifact of the software tool or a result of smORF translation being generally noisier and more difficult to assess relative to larger annotated coding regions. To test this, we initially examined a high-resolution HeLa-S3 Ribo-seq data for differences in identifying translated ORFs across the different tools. We observed high overlap in the number of total annotated genes detected across all five tools with 8,781 (71.6%) called translated and a similar number identified by each tool (**Fig. 2A**). Pairwise comparisons of the number of annotated ORFs found in one tool compared to each other tool, as well as the proportion of matched ORFs, showed similar performance between all tools and that RibORFv0.1 was the least sensitive (**Figs. 2B, C**). Next, we examined the prediction of novel translated smORFs from each tool (**Fig. 2D**). Compared to annotated ORFs, there is little overlap in the total number of smORFs predicted with only 235 (2.3%) found across all tools and 1,549 (15.4%) smORFs found in at least three out of five tools. The performance of the tools differentiated into two groups. RiboCode, RibORFv0.1, and RibORFv1.0 called 2.3-4.8 times as many smORFs translated as ORFquant and Ribo-TISH. Pairwise analysis of the number and proportion of matched smORFs revealed additional differences between the tools (**Fig. 2E, F**). First, despite identifying less than half the number of translated smORFs as RiboCode and RibORFv1.0, only ∼40% of Ribo-TISH hits overlapped with RiboCode and RibORFv1.0. This contrasted with ORFquant, which also identified a lower amount of translated smORFs (1,124) but had 68% and 81% of its calls overlap with those of RibORFv1.0 and RiboCode, respectively. In addition, Ribo-TISH had the smallest proportion of ORFquant calls matched (30%). These data demonstrate that Ribo-TISH is an outlier compared to the other tools that identifies both a smaller number and more unique set of smORFs as translated. Meanwhile, the majority of ORFquant’s hits can be captured by using the tools that predict larger numbers of translated smORFs.

**Figure 2.**
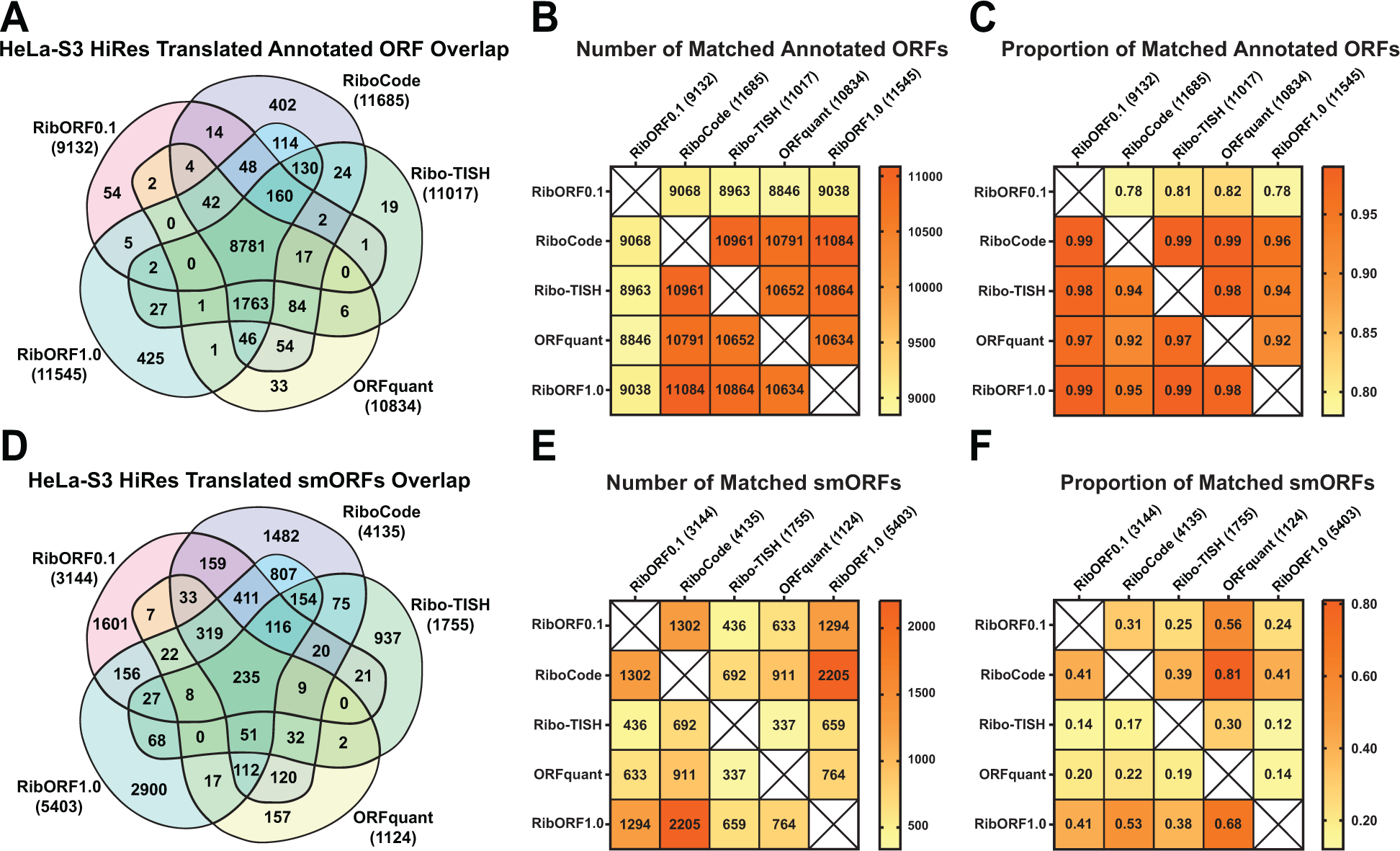
Comparison of detected annotated ORFs and predicted smORFs in the high-resolution HeLa-S3 Ribo-seq dataset. (A) Venn diagram showing the overlap of annotated genes called translated across the different tools. The total number of annotated genes detected is displayed next to the names of each tool in parentheses. **(B)** Heat map showing the pairwise comparison of matching annotated genes between the different tools. **(C)** Heat map showing the proportion of annotated genes identified by the tool in the column that are also detected by the tool in the row. **(D-E)** The same plots are shown as in **(A-C)** for the analysis of unannotated smORFs.

We next explored whether these trends would remain consistent after analyzing low-resolution HeLa-S3 Ribo-seq data. Compared to the detection of annotated genes in the high-resolution dataset, we observed a large drop in the number of smORFs called translated by ORFquant (3,525) and Ribo-TISH (5,894) resulting in only 2,104 (18.4%) in common across all tools (Fig. 3A). Pairwise comparisons of the tools showed that both RibORFv1.0 and RiboCode identified the most annotated genes as translated and >90% of those identified in all the other tools (Fig. 3B, C). ORFquant was impacted the most by the low-resolution data, identifying only 3,525 annotated genes as translated. This is consistent with ORFquant’s requirement to have >50% reads in-frame for each codon within an ORF to be called translated [9]. Similarly, Ribo-TISH and ORFquant were greatly affected by the lower resolution when predicting novel smORFs, predicting 13 and 203 smORFs, respectively (Fig. 3D). Despite the low number of smORFs predicted, we observed the same trend that the highest proportional overlap of smORFs were relative to ORFquant predictions. We repeated the comparison analysis on HEK293T Ribo-seq data with varying resolutions and found similar trends with annotated genes (**Supplementary** Fig. 3) and unannotated smORFs (**Supplementary** Fig. 4), validating the conclusion that ORFquant and Ribo-TISH are less noise tolerant for calling translation of both annotated genes and translated smORFs compared to the other three pipelines.

**Figure 3.**
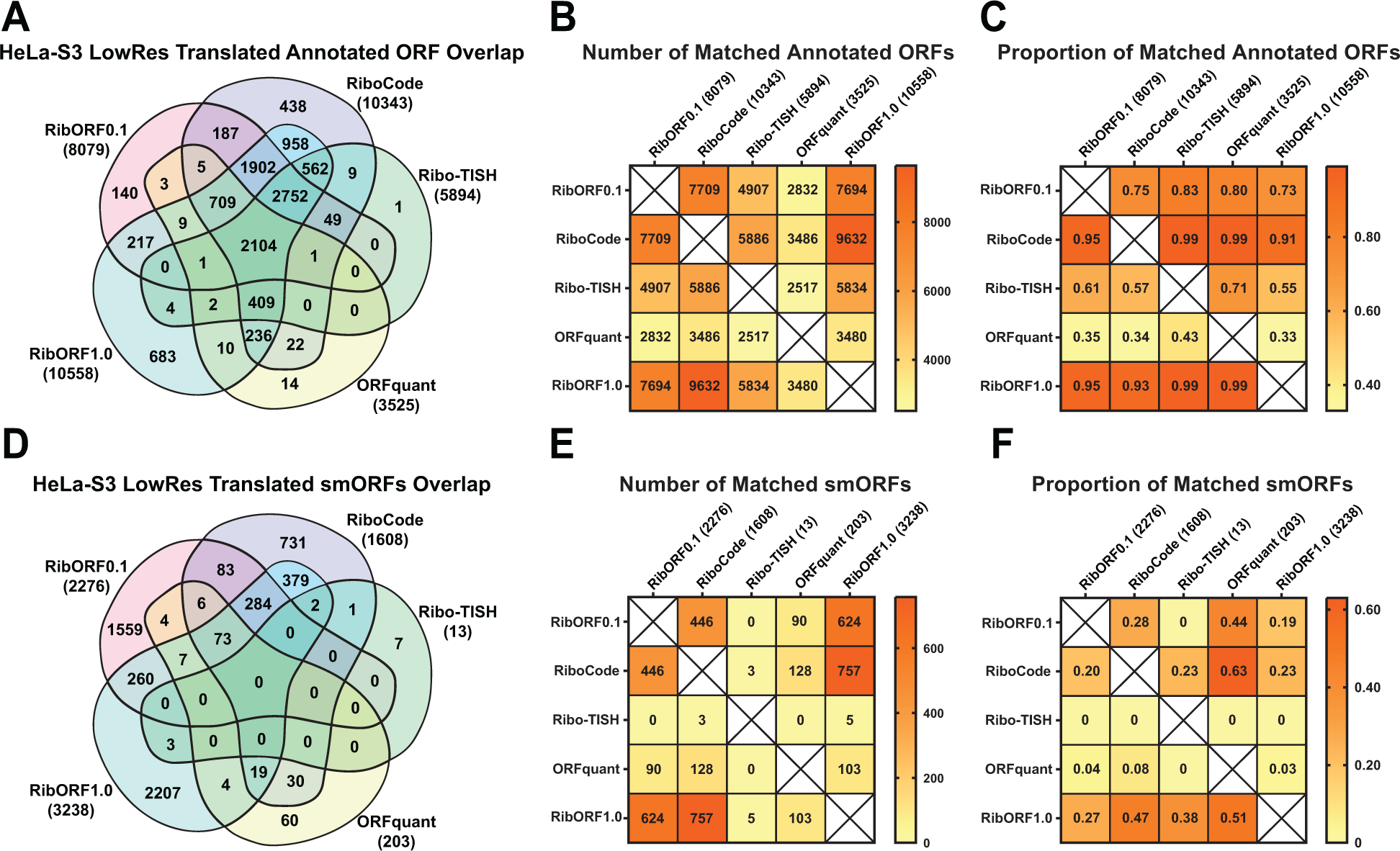
Comparison of detected annotated ORFs and predicted smORFs in low-resolution HeLa-S3 Ribo-seq dataset. (A) Venn diagram showing the overlap of annotated genes called translated across the different tools. The total number of annotated genes detected is displayed next to the names of each tool in parentheses. **(B)** Heat map showing the pairwise comparison of matching annotated genes between the different tools. **(C)** Heat map showing the proportion of annotated genes identified by the tool in the column that are also detected by the tool in the row. **(D-E)** The same plots are shown as in **(A-C)** for the analysis of unannotated smORFs.

We also directly compared smORF predictions across the low- and high-resolution HeLa-S3 datasets (**Supplementary** Fig. 5). Despite using lower quality Ribo-seq data, both versions of RibORF and RiboCode were all able to identify a small fraction of smORFs (between 10-25%) that were also called translated by each of the five tools when using high resolution data. These results suggest that RibORF and RiboCode can identify smORFs that are highly likely to be translated despite the use of lower resolution data, though many of the smORFs called translated are still likely to be noise. For ORFquant, only 1-5% of ORFs called translated using low-resolution dataset were also observed by other tools when using high resolution data, while Ribo-TISH only identified 1-3 total hits in common when comparing low versus high resolution data. Overall, RibORF and RiboCode demonstrate more sensitive detection of translated smORFs than ORFquant and Ribo-TISH regardless of data quality.

### Accounting for Isoform Differences in smORF Predictions

In our initial comparisons between the tools, we restricted the matches to smORFs that have the same genomic coordinates. However, given that smORFs can use alternative start codons and can be spliced like larger ORFs, it is possible that the tools predict isoforms of the same smORF. To account for this, we looked for any additional smORFs identified by each tool that have the same start coordinate but different stop coordinates and vice versa using our HeLa-S3 datasets. Each tool was pairwise compared against RibORFv1.0, which predicted the largest number of smORFs. For the high-resolution dataset, allowing for stop site matches (start site isoforms) resulted in an additional 79 to 411 smORFs in common, while allowing for start site matches (stop site isoforms) resulted in an additional 6 to 48 smORFs in common (**Fig. 4A**). The high number of start site isoforms called between the different tools is expected due to how the different pipelines handle AUG versus near cognate start codons as well as ORFs where multiple possible start codons are present. For example, ORFquant will only allow for AUG start codons in its predictions. For the low-resolution HeLa-S3 dataset, additional matching smORFs were found for RibORF and RiboCode, but very few additional hits were observed for ORFquant and Ribo-TISH due to the overall lower number of smORFs called translated by these tools (**Fig. 4B**). Examples of predicted smORF isoforms that have matched start or stop coordinates can be observed in the 5’-UTRs of CES and ANGEL2, respectively (**Fig. 4C**).

**Figure 4.**
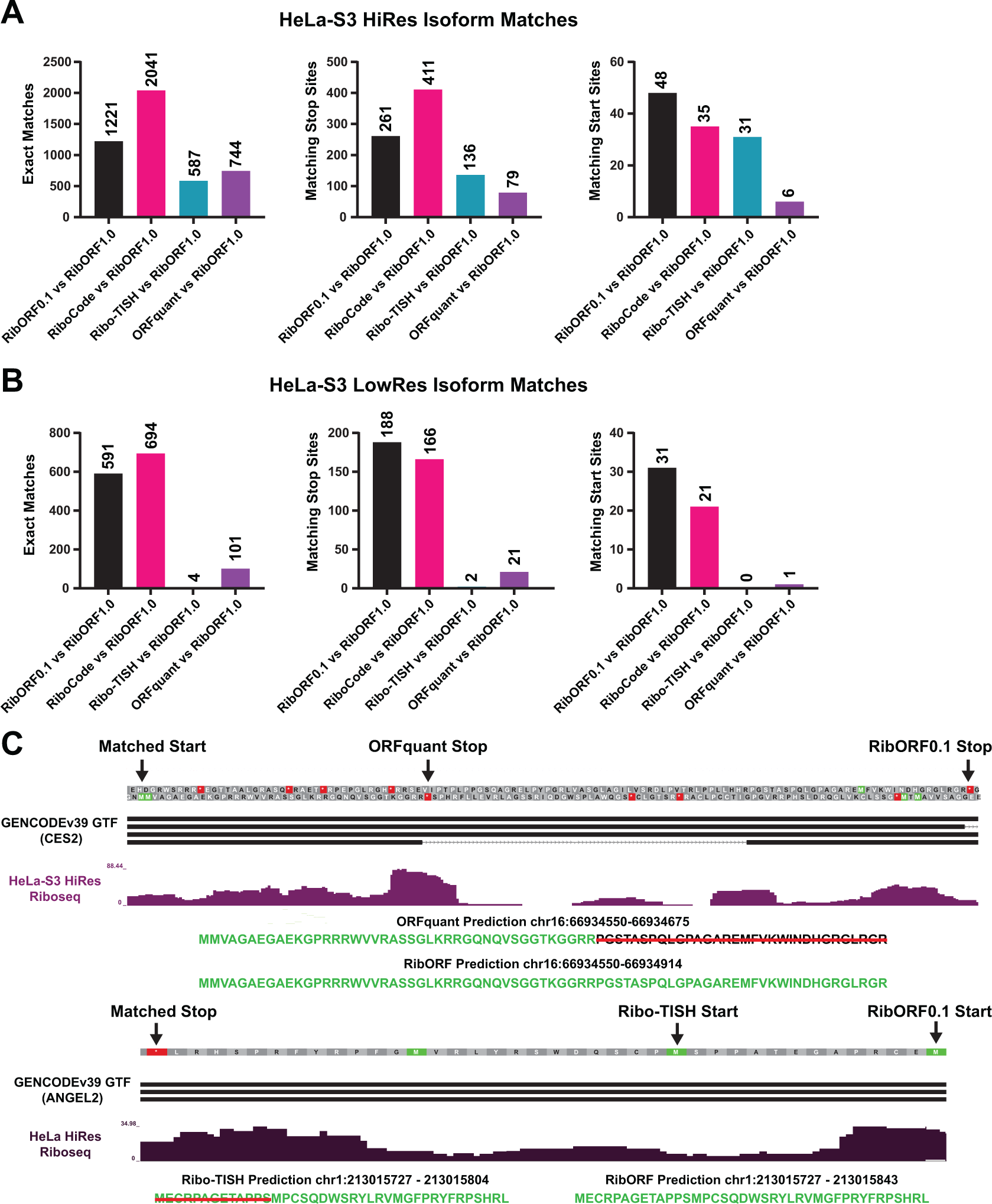
Accounting for smORF isoform variance across different ORF prediction pipelines. (A-B) Bar plots showing the number of exact smORF matches (left), start site isoform smORF matches (middle), and stop site isoform smORF matches (right) between each tool and RibORFv1.0 when analyzing either the high-resolution **(A)** or low-resolution **(B)** HeLaS3 Ribo-seq datasets. **(C)** Bedgraph tracks showing Ribo-seq coverage on the 5’-UTRs of CES2 and ANGEL2. In the top track, an alternatively spliced smORF on the positive strand was identified by both ORFquant and RibORFv0.1 with a matching start site but different the stop site. In the bottom track, Ribo-TISH and RibORFv0.1 detect a smORF on the negative strand with the same stop location but different canonical start codons.

### Incorporating Translation Initiation Sequencing (TI-Seq) Data into smORF Prediction

To aid in the prediction of novel ORFs, some newer tools like Ribo-TISH allow integration of translation initiation sequencing (TI-seq). TI-seq is a modified version of Ribo-seq that includes a short pretreatment with translation initiation inhibitors such as harringtonin or lactimidomycin in order to enrich for ribosome coverage on ORF start sites, providing additional evidence of their translation [12]. Using matched TI-seq HeLa-S3 data from harringontin treated cells, we compared annotated genes and smORFs called translated when using both TI-seq and standard Ribo-seq datasets to those identified by Ribo-seq alone. There was a high overlap of annotated genes detected (∼73%), though fewer total genes were called translated when TI-seq data was included due to the extra requirement of having an initiation peak (**Supplementary** Fig. 6A). For smORFs, the overlap between the two analyses was much lower (∼10%, **Supplementary** Fig. 6B). In some instances, the lack of overlap was due to different translation start sites predicted based on whether TI-seq data was incorporated or not. We highlight one example of two smORF isoforms on the TXNRD1 transcript, with one smORF starting at an AUG start codon that shows enrichment by TI-seq and the other starting at an upstream near cognate start codon that is predicted when using Ribo-seq alone (**Supplementary** Fig. 6C). While differing start site predictions can explain some of the differences, some of the smORFs identified by Ribo-seq alone using Ribo-TISH might in fact not be translated since they did not show start site enrichment by TI-seq. Ribo-TISH also predicts unique smORFs found only with integration of initiation site data, such as the smORF within the 5’-UTR of the PIGW transcript (**Supplementary** Fig. 6D). Thus, the inclusion of initiation site data can introduce another variable to smORF predictions.

### Impact of *de novo* Assembled Transcriptome Annotation on smORF Identification

Analyzing Ribo-seq data for translated smORFs requires the use of a transcriptome to create a database of all possible smORFs present in a given sample. While most studies use transcriptomes sourced from reference databases like GENCODE [13] or Ensembl [14], *de novo* assembled transcriptomes can also be used. By incorporating *de novo* transcriptome assemblies, one can identify smORFs on transcript isoforms that are otherwise missing from these public reference databases. We previously used transcriptomes assembled from Illumina-based short read RNA-seq data to identify smORFs on cell line specific transcript isoforms [3], but use of long-read sequencing technologies may aid in the identification of additional smORFs. To evaluate the two sequencing methods’ effects on smORF identification, we assembled HeLa-S3 transcriptomes from both Nanopore long-read and Illumina short-read

RNA-seq datasets using StringTie [15], a more modern assembly tool than what we had used in our original study. After assembly, the resulting transcriptome was merged with the GENCODE reference to create a comprehensive transcriptome that includes additional transcripts identified by each RNA-seq strategy. This resulted in an additional 40 transcripts using Illumina RNA-seq data and an additional 1,106 transcripts using Nanopore RNA-seq data that were not included in the GENCODE transcriptome (**Fig. 5A**). Using RibORFv0.1 to identify translated smORFs in the high-resolution HeLaS3 dataset with each *de novo* assembled transcriptome revealed a high degree of overlap (∼94%, **Fig. 5B**). However, unique predicted translated smORFs were found for each transcriptome, with 127 predicted smORFs found only when using the Nanopore assembly and 69 specifically from the Illumina assembly. Using RiboCode for translation calling yielded similar results (**Supplementary** Fig. 7). An example smORF that both RibORFv0.1 and RiboCode call translated from a transcript specifically identified when using Nanopore long-read RNA-seq data can be found antisense to ADARB2 (**Fig. 5C**). These data show that incorporating *de novo* transcriptome assembly into smORF prediction workflows can identify additional hits, but the overall benefit over using the GENCODE reference transcriptome alone is marginal.

**Figure 5:**
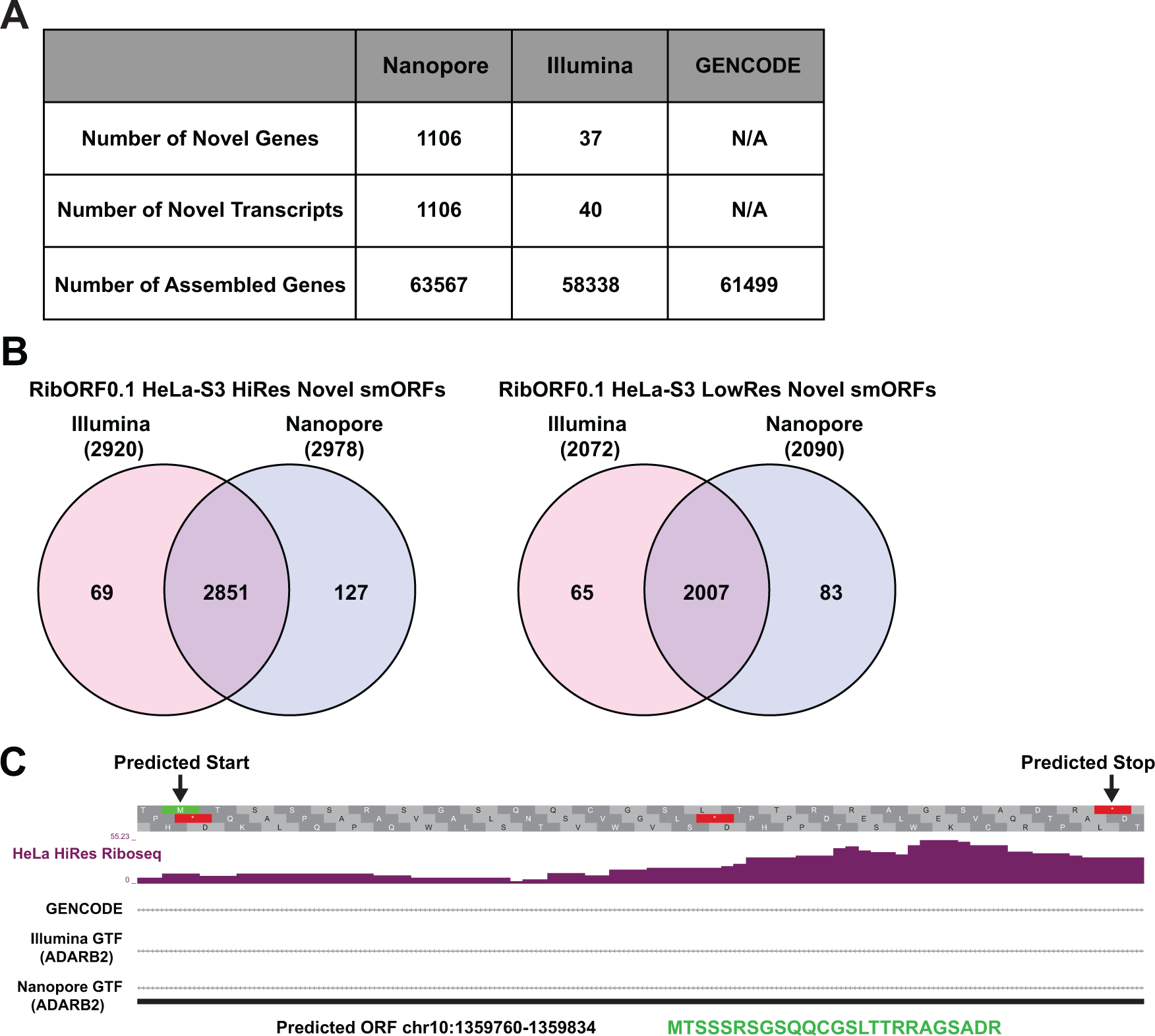
*De novo* transcriptome assembly enables additional smORFs to be predicted. **(A)** Comparison of the Nanopore long read- and Illumina short read-based *de novo* assembled transcriptomes and GENCODE reference using GffCompare. **(B)** Venn diagram showing the overlap of predicted smORFs identified by RibORFv0.1 when using the de novo transcriptome assemblies along either high-resolution (left) or low-resolution (right) HeLa-S3 Ribo-seq datasets. The total number of annotated genes detected is using each assembly is shown in parentheses. **(C)** Bedgraph tracks showing Ribo-seq coverage on a region antisense to ADARB2 as well as transcripts present in GENCODE and the de novo transcriptome assembles. An assembled transcript for this region is only found when using the Nanopore-based *de novo* assembly.

### Comparing Tool Accuracy with a High Confidence Community smORF Dataset

Comparing predicted translated smORFs across tools showed high variability, leading one to question which tool is better at identifying *bona fide* microprotein-coding smORFs. To address this point, we compared the predicted smORFs from each tool to a community set of 3,085 smORFs from different human samples that were reproducibly detected across multiple Ribo-seq-based smORF annotation studies and using different tools [16]. These high confidence smORF annotations are publicly available through GENCODE. Using the HeLa-S3 datasets, we determined the number of smORFs matching the GENCODE smORF set for each tool. For the high-resolution HeLa-S3 dataset, 155 of these high confidence GENCODE smORFs were predicted by all tools, and each was able to identify a subset of these smORFs missed by the other tools (**Fig. 6A**). RibORFv0.1, RibORFv1.0, and RiboCode had the highest number of matches, consistent with their overall greater number of smORFs called translated compared to Ribo-TISH and ORFquant (**Fig. 6B**). However, ORFquant had the highest proportion of its smORF calls overlap with the GENCODE set. Similar trends are observed when using the low-resolution HeLa-S3 dataset, with the exception that Ribo-TISH and ORFquant call far fewer smORFs than the other tools when using poorer quality data (**Supplementary** Fig. 8). Overall, these results further demonstrate that RibORF and RiboCode are more sensitive than ORFquant, while ORFquant is the most accurate of the tools and Ribo-TISH suffers from both lower sensitivity and accuracy.

**Figure 6.**
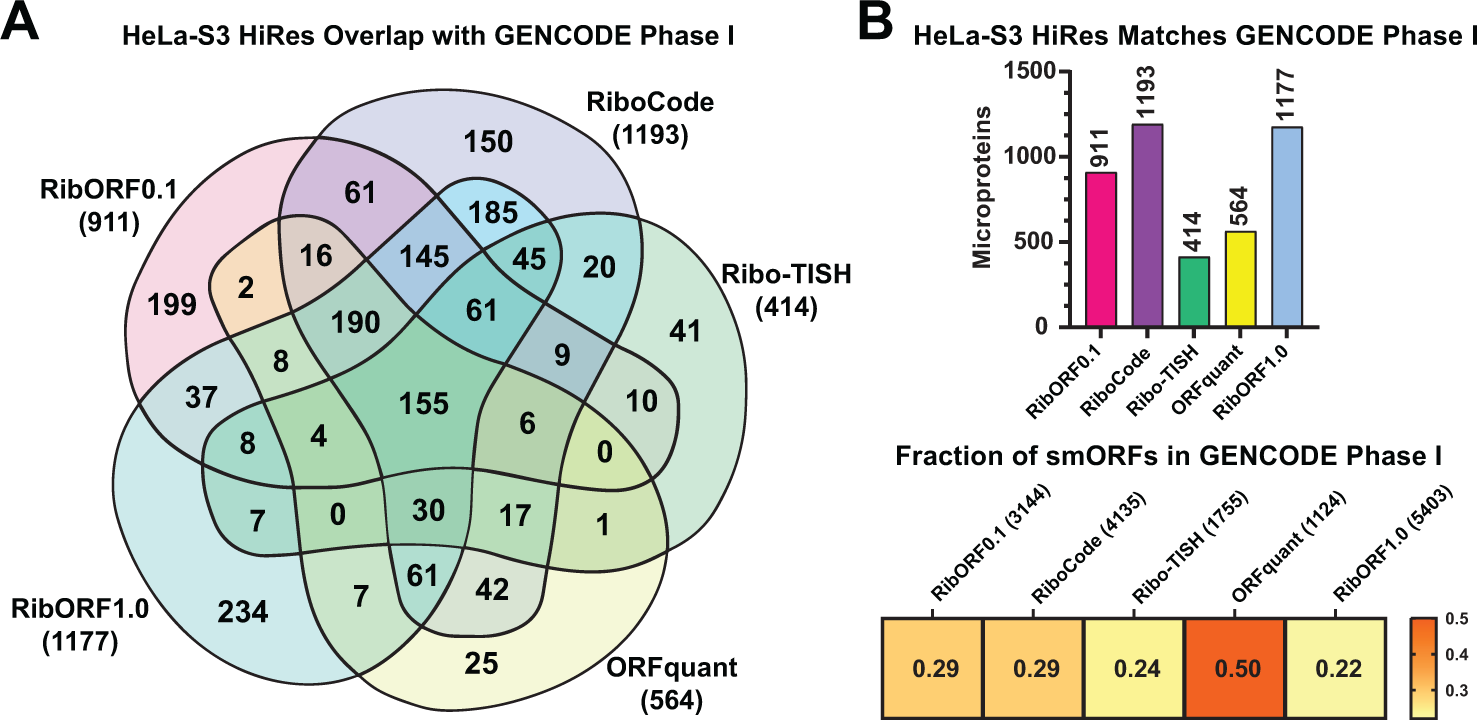
Comparison of the GENCODE Phase I high-confidence smORFs predicted by each tool in the HeLa-S3 high-resolution dataset. (A) Venn diagram showing the overlap of GENCODE smORFs detected by each tool in the high-resolution HeLaS3 Ribo-seq dataset. Total number of GENCODE smORFs detected by each tool is shown in parentheses. **(B)** Bar plot showing the total number of matched smORFs with the GENCODE set for each tool (top). Heat map showing the proportion of smORFs identified by each tool that are also included in the GENCODE smORF set (bottom).

### Translation Levels Correlate with smORF Detectability by Multiple Tools

Given the high overlap of annotated genes called translated across the different tools but low overlap of predicted translated smORFs, we wanted to identify properties that influence this difference. Ribo-seq read coverage for a given ORF correlates with translation levels and is a critical factor in predicting translation for each of these tools. Therefore, we compared Ribo-seq read coverage of annotated genes and smORFs called translated by each tool. Using the high-resolution HeLaS3 dataset, annotated genes called translated showed significantly higher in all tools except Ribo-TISH (**Supplementary** Fig. 9). These same patterns were observed when analyzing the low-resolution Ribo-seq dataset using RiboCode and RibORF (**Supplementary** Fig. 10). These data suggest that overall higher translation levels are likely driving the greater overlap in annotated gene detection across the different tools. We therefore hypothesized that smORFs that are reproducibly detected across the different tools are also more likely to have higher translation levels. Comparing smORFs called translated by all five, at least three, and less than three tools showed that smORFs detected by more tools are translated at significantly higher levels in both high- and low-resolution datasets (**Supplementary** Fig. 11A). These results suggest that smORFs are more difficult to detect in part because of their overall lower translation levels than larger annotated ORFs.

Most human microprotein-coding smORFs show conservation only to other primates or are entirely de novo occurrences in our genome [17–19]. However, there are many examples of functionally characterized microproteins that are well conserved across mammals [20,21].

Therefore, we next assessed whether smORFs detected by multiple tools are not only translated at higher levels but also more conserved. Using PhyloCSF [22], we observed no significant difference in average scores between smORFs detected by three or more tools and those detected by fewer than three tools (**Supplementary** Figure 11B). Thus, conservation is not a major determinant of high confidence smORF detection by Ribo-seq.

## CONCLUSIONS

Ribo-seq has revolutionized our ability to *de novo* annotate translated open reading frames. Still, it is only as effective as the bioinformatic tools used to interpret the data to identify *bone fide* translation events. By comparing several popular tools, we found that each can identify a similar set of translated annotated genes as intended when high-resolution data is used. When attempting to identify unannotated translated smORFs, however, the tools vary widely in the number called translated and show little overlap. We found a clear split between RibORFv0.1, RibORFv1.0, and RiboCode, which consistently predict more translated smORFs than ORFquant and Ribo-TISH. Moreover, RiboCode and RibORFv1.0 identify a large fraction of the same smORFs called by ORFquant, while Ribo-TISH identifies a subset of smORFs that is more unique than all the other tools. When low-resolution Ribo-seq data is used, ORFquant and Ribo-TISH are further separated from the other tools, identifying a relatively small number smORFs as translated and reflecting differences in stringency. When comparing the smORFs predicted by each tool with a high confidence set included in GENCODE, we found that RiboCode and RibORF had the highest sensitivity but ORFquant the highest accuracy. Given these results, we suggest that RiboCode and both versions of RibORF are better suited for identifying smORFs to test in high-throughput screens like CRISPR dropout assays where the aim is to identify large sets of functional smORFs. These tools are also good choices when only lower quality Ribo-seq data is available, though caution must be exercised as lower-resolution data will inherently lead to noisier calls overall. ORFquant, meanwhile, is an excellent choice when attempting to identify confidently translated smORFs with AUG start sites from high-resolution data, as when planning low-throughput functional characterization studies of encoded microproteins. Finally, we suggest that regardless of the purpose it is prudent to use multiple Ribo-seq analysis tools in addition to analyzing biological replicates to identify the most confident microprotein-coding smORFs, particularly for ongoing annotation efforts for reference databases, and consider Ribo-seq read coverage in their prioritization.

## ACKNOWLEDGEMENTS

We thank members of the Martinez lab for thoughtful suggestions and comments throughout the study. We also thank Dr. Jorge Ruiz Orera for helpful advice on running ORFquant.

## AUTHOR CONTRIBUTIONS

G.T. and T.F.M. conducted the computational experiments and analyzed the results; N.H. prepared Nanopore long-read RNA-seq data; T.F.M. conceived, organized, and managed the project implementation; all authors wrote and reviewed the manuscript.

## SUPPLEMENTARY DATA

**Supplementary Figure 1.**
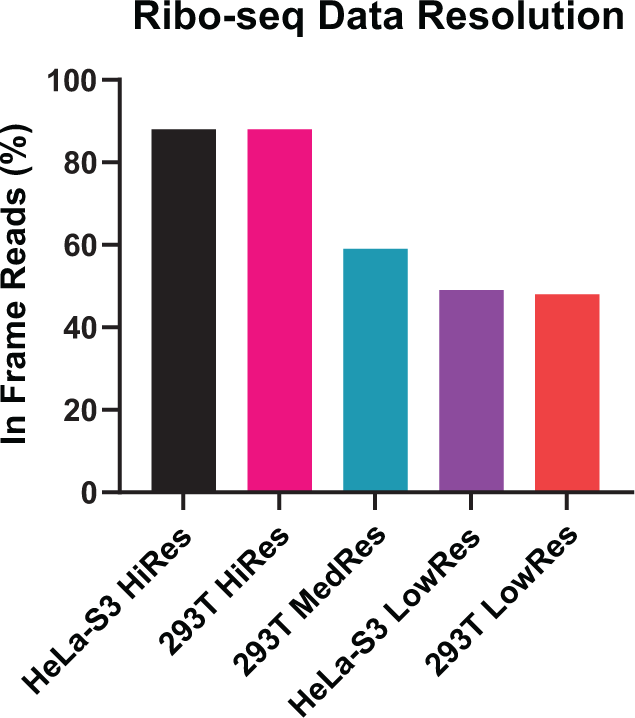
Percentage of in-frame reads for the highest abundance RPF read length for each Ribo-seq dataset. For each Ribo-seq dataset analyzed, the fraction of in-frame reads after the start site was calculated for the most abundant RPF read length after offset correction to align to the ribosomal A-site. The read lengths analyzed for each dataset were: HeLa-S3 HiRes - 28 nt, 293T HiRes - 28 nt, 293T MedRes - 30 nt, HeLa-S3 LowRes – 32 nt, and 293T LowRes - 31 nt.

**Supplementary Figure 2.**
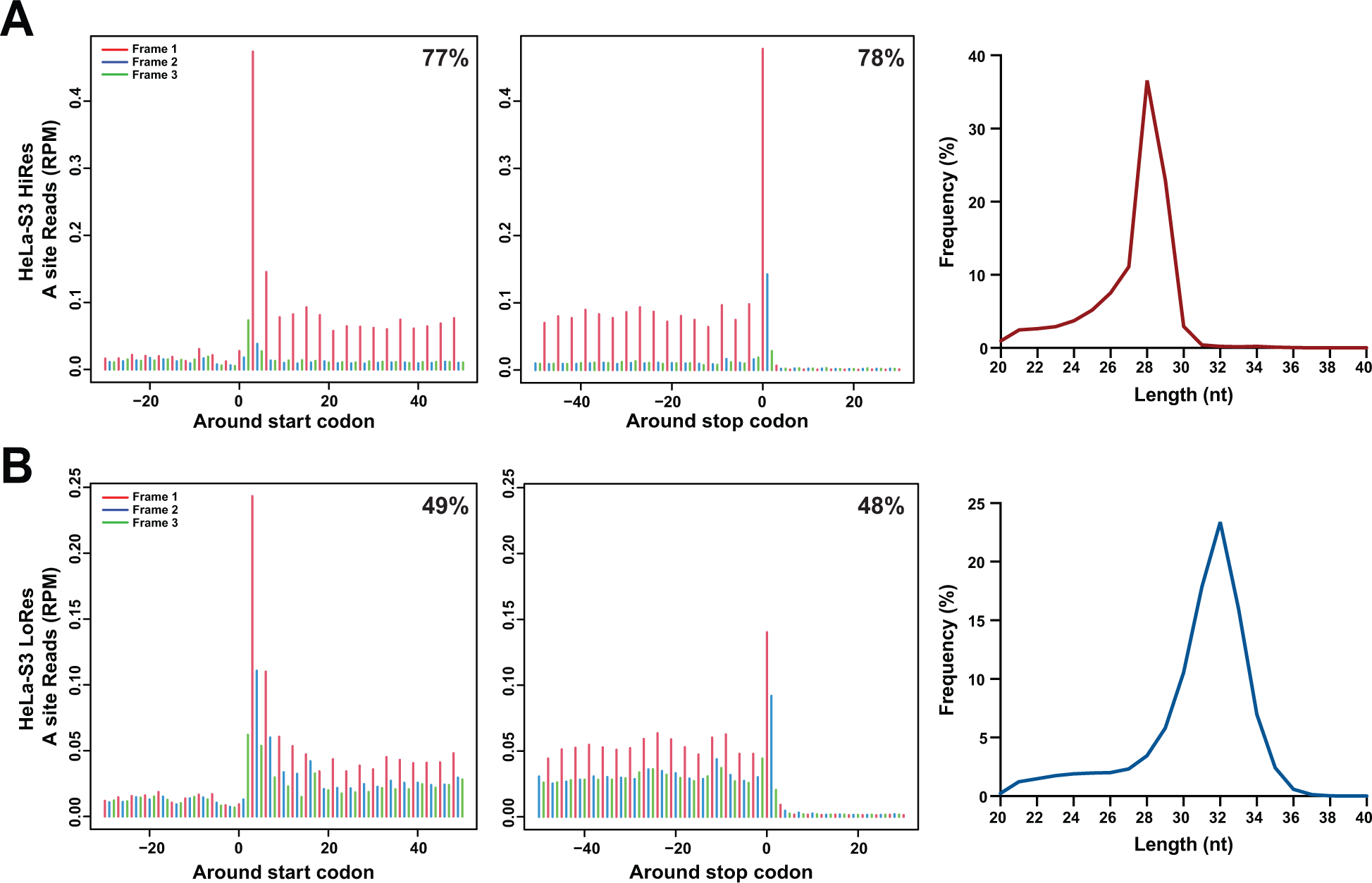
Quality control of HeLa-S3 Ribo-seq datasets. (A) Metagene plots of the high-resolution HeLa-S3 dataset displaying the A-site read distribution around the start and stop sites (left). The 5’-end of each RPF read was adjusted to the ribosomal A-site after mapping to hg38 canonical genes. The coding regions are in reading frame 1, while reading frames 2 and 3 are out of frame. Line graph of the RPF read length frequency distribution peaks at 28 nt (right). Read lengths 25-29 nt were used for downstream analysis **(B)** Metagene plots of the low-resolution HeLa-S3 dataset displaying the A-site read distribution around the start and stop sites (left). Line graph of the RPF read length frequency distribution peaks at 32 nt. Read lengths 31-35 nt were used for downstream analysis.

**Supplementary Figure 3.**
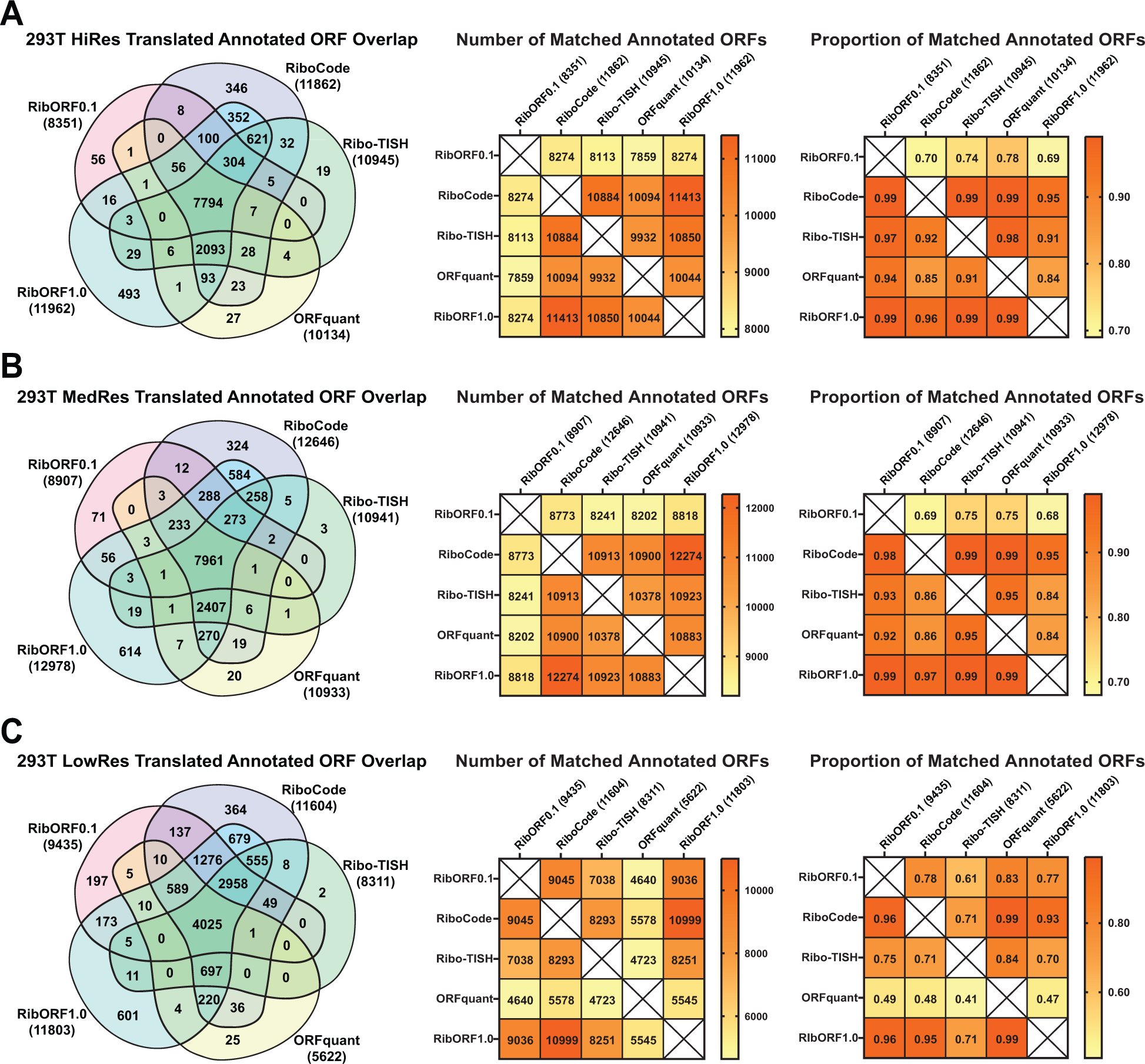
Comparison of detected annotated ORFs in HEK293T Ribo-seq datasets of varying resolution. (A-C) Venn diagram showing the overlap of annotated genes called translated across the different tools (left). The total number of annotated genes detected is displayed next to the names of each tool. Heat map showing the pairwise comparison of matching annotated genes between the different tools (middle). Heat map showing the proportion of annotated genes identified by the tool in the column that are also detected by the tool in the row (right). HEK293T datasets analyzed are categorized by their resolution: high **(A)**, medium **(B)**, and low **(C)**.

**Supplementary Figure 4.**
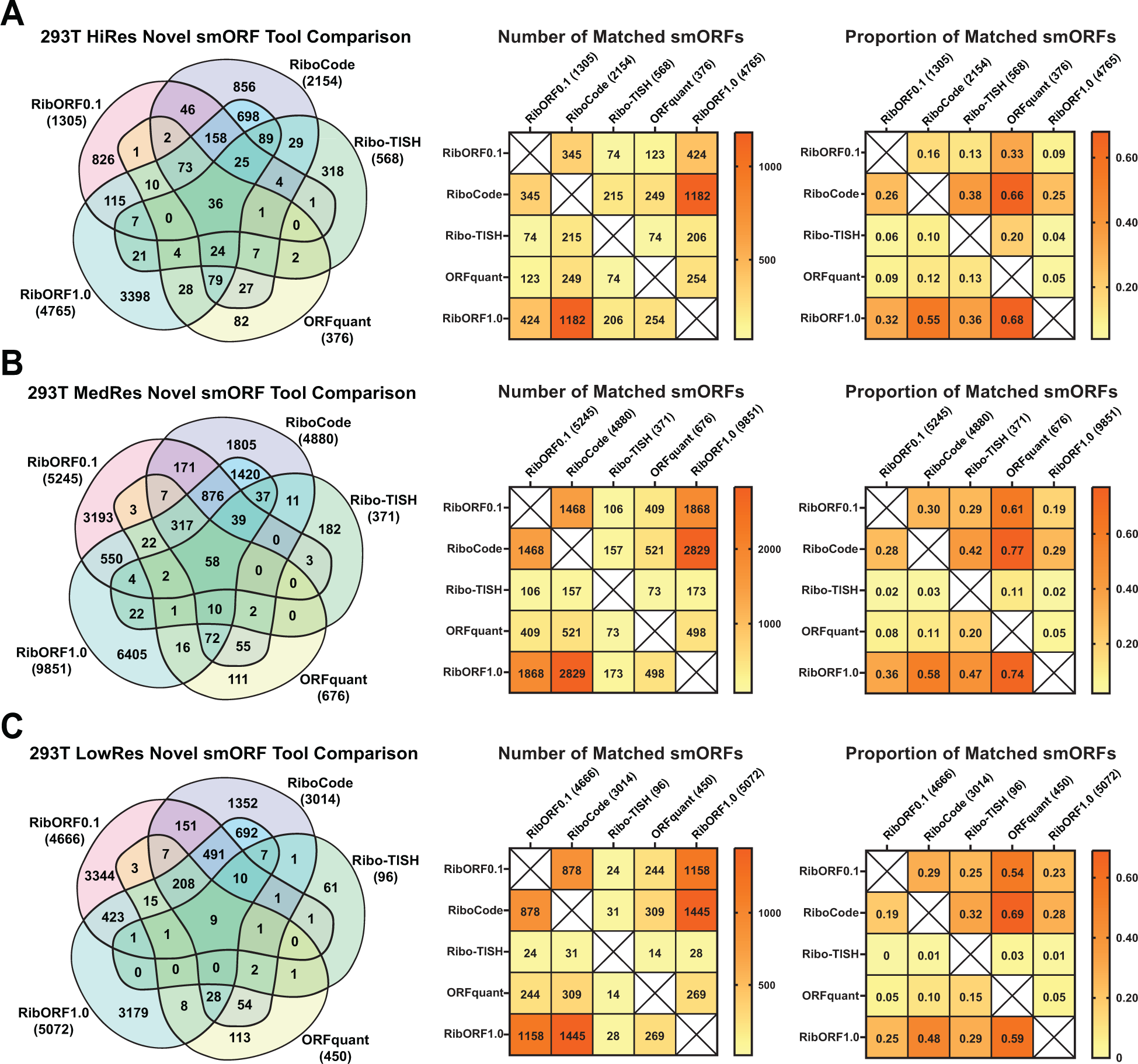
Comparison of predicted smORFs in HEK293T Ribo-seq datasets of varying resolution. (A-C) Venn diagram showing the overlap of unannotated smORFs called translated across the different tools (left). The total number of smORFs detected is displayed next to the names of each tool. Heat map showing the pairwise comparison of matching annotated genes between the different tools (middle). Heat map showing the proportion of unannotated smORFs identified by the tool in the column that are also detected by the tool in the row (right). HEK293T datasets analyzed are categorized by their resolution: high **(A)**, medium **(B)**, and low **(C)**.

**Supplementary Figure 5.**
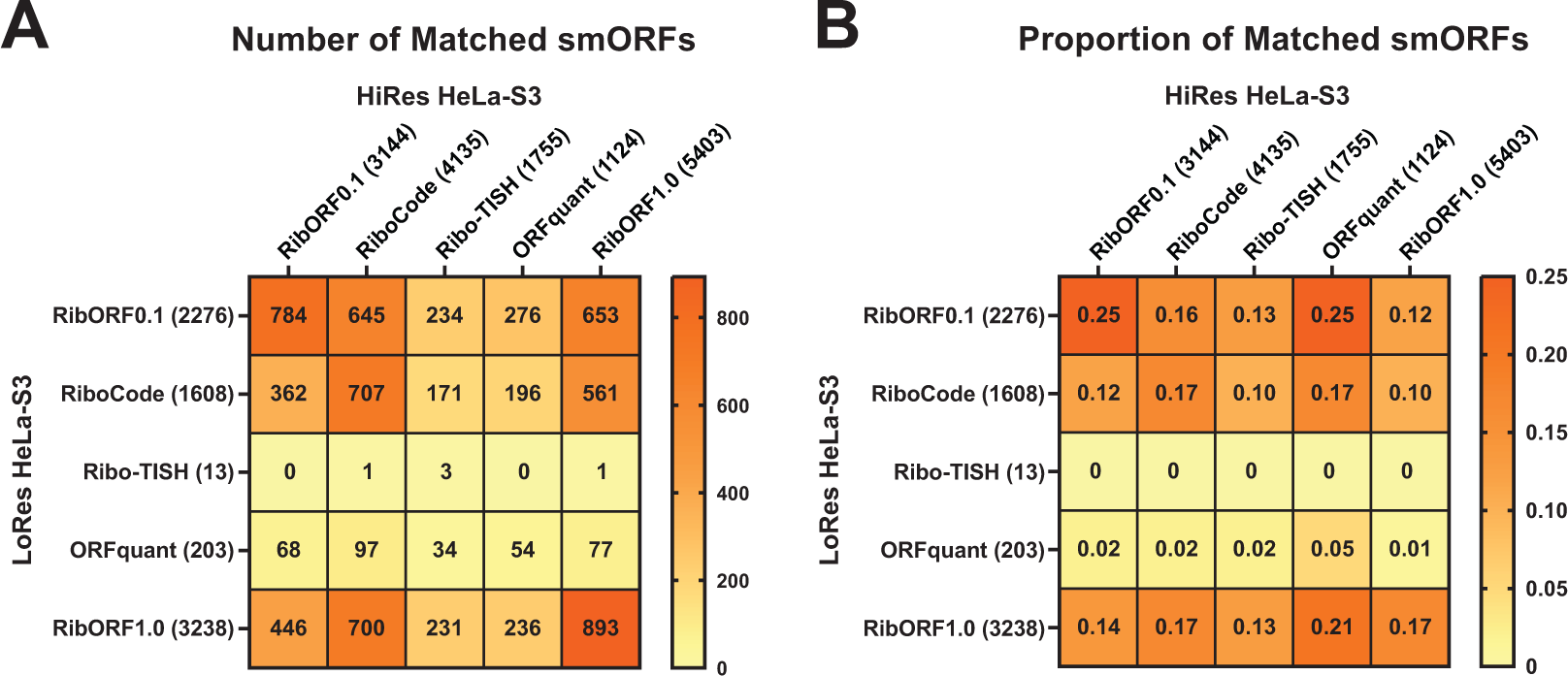
Comparison of predicted smORFs from high- and low-resolution HeLa-S3 Ribo-seq datasets. (A) Heat map showing the pairwise comparison of matching smORFs predicted in both high- and low-resolution HeLa-S3 Ribo-seq datasets. **(B)** Heat map showing the proportion of unannotated smORFs identified by the tool in the column using high-resolution data that are also detected by the tool in the row using low-resolution data. The total number of smORFs detected by each tool is shown in parentheses.

**Supplementary Figure 6.**
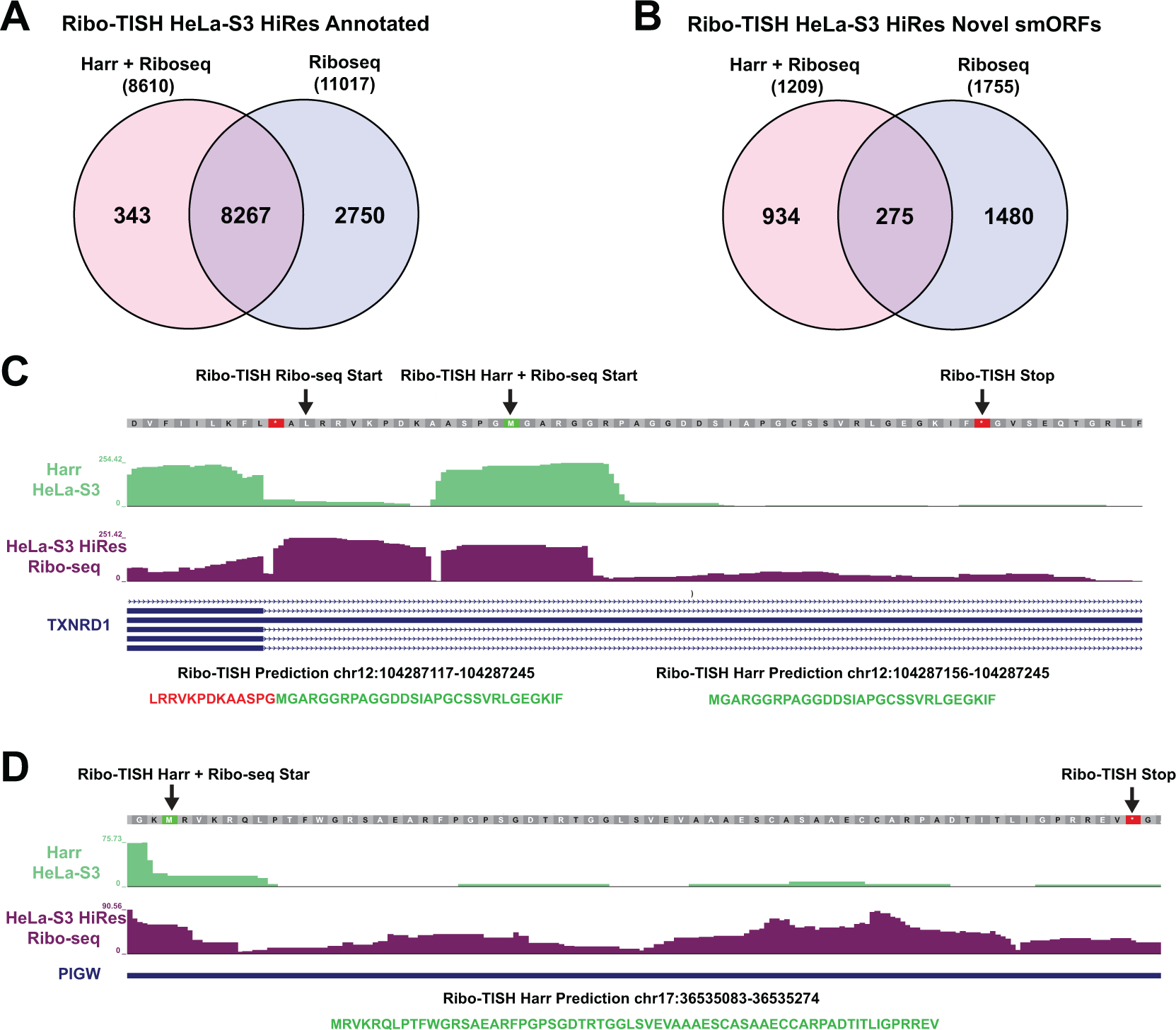
Comparison of predicted smORFs using Ribo-TISH with translation initiation (TI-seq) data included and Ribo-seq data alone. (A) Venn diagram showing the overlap of annotated ORFs detected by Ribo-TISH including or excluding TI-seq HeLa-S3 data along with high-resolution Ribo-seq data. Total number of annotated ORFs identified is displayed next to each condition analyzed in parentheses. **(B)** Same analysis as in **(A)** for predicted smORFs. **(C)** Bedgraph tracks showing TI-seq and Ribo-seq coverage for smORFs called translated within the 5’-UTR of the TXNRD1 transcript on the positive strand. Both smORFs share the same stop site but have different starts called if TI-seq data is considered. (**D)** Bedgraph tracks showing a smORF identified within the 5’-UTR of PIGW only when running Ribo-TISH with both TI-seq and Ribo-seq data.

**Supplementary Figure 7.**
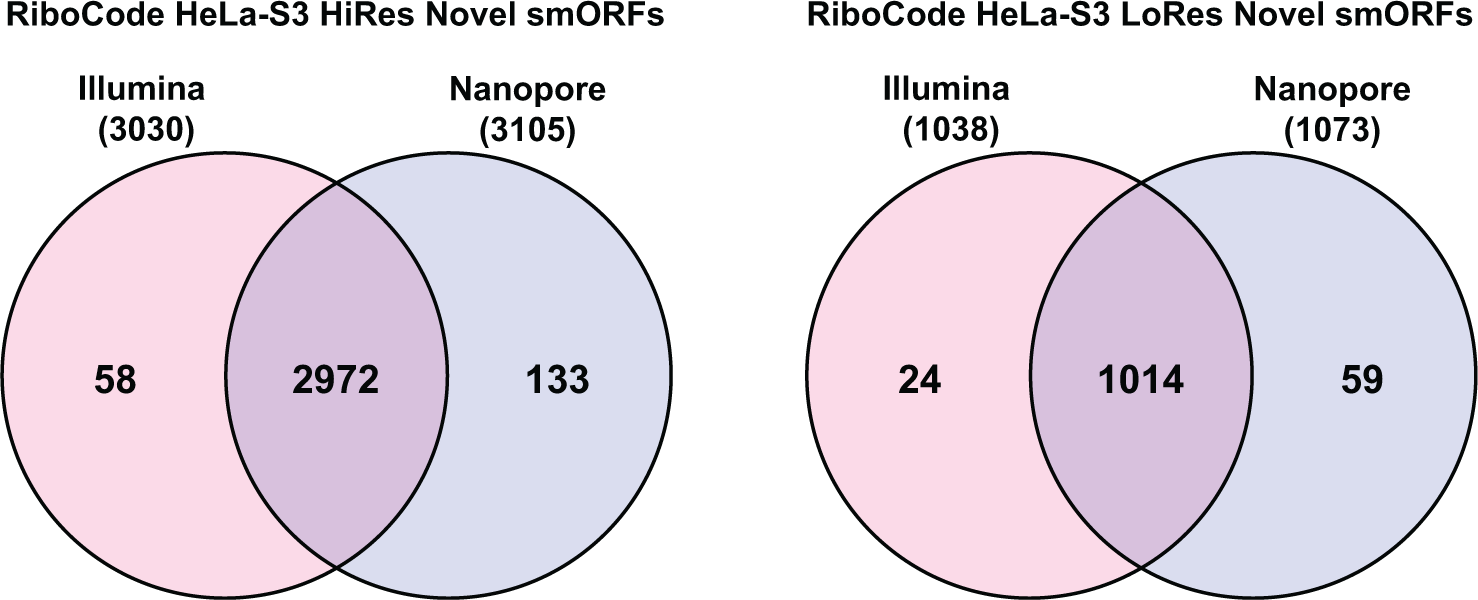
Comparison of smORFs predicted by RiboCode when incorporating Nanopore- and Illumina-based *de novo* assembled transcriptomes. Venn diagram showing the overlap of predicted smORFs identified by RiboCode when using the *de novo transcriptome* assemblies along either high-resolution (left) or low-resolution (right) HeLa- S3 Ribo-seq datasets.

**Supplementary Figure 8.**
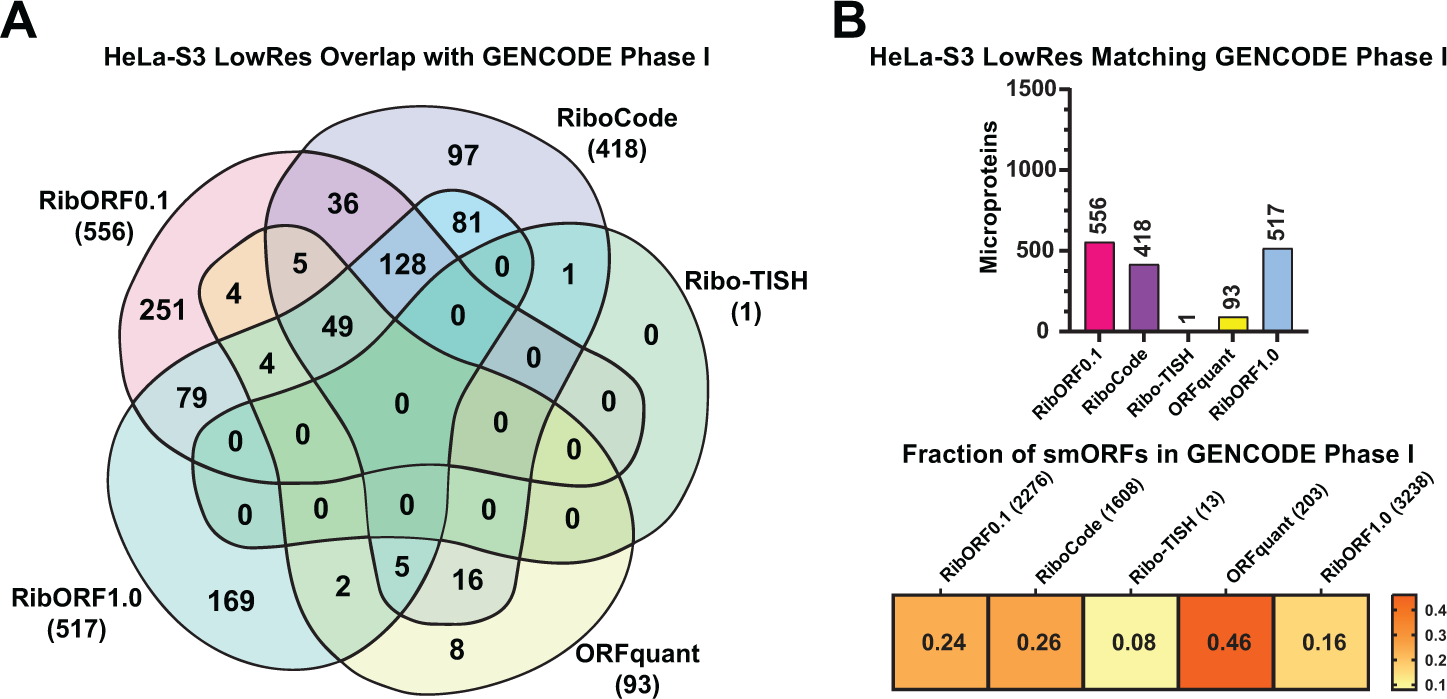
Comparison of the GENCODE Phase I high-confidence smORFs predicted by each tool in the HeLa-S3 low-resolution dataset. (A) Venn diagram showing the overlap of GENCODE smORFs detected by each tool in the low-resolution HeLaS3 Ribo-seq dataset. Total number of GENCODE smORFs detected by each tool is shown in parentheses. **(B)** Bar plot showing the total number of matched smORFs with the GENCODE set for each tool (top). Heat map showing the proportion of smORFs identified by each tool that are also included in the GENCODE smORF set (bottom).

**Supplementary Figure 9.**
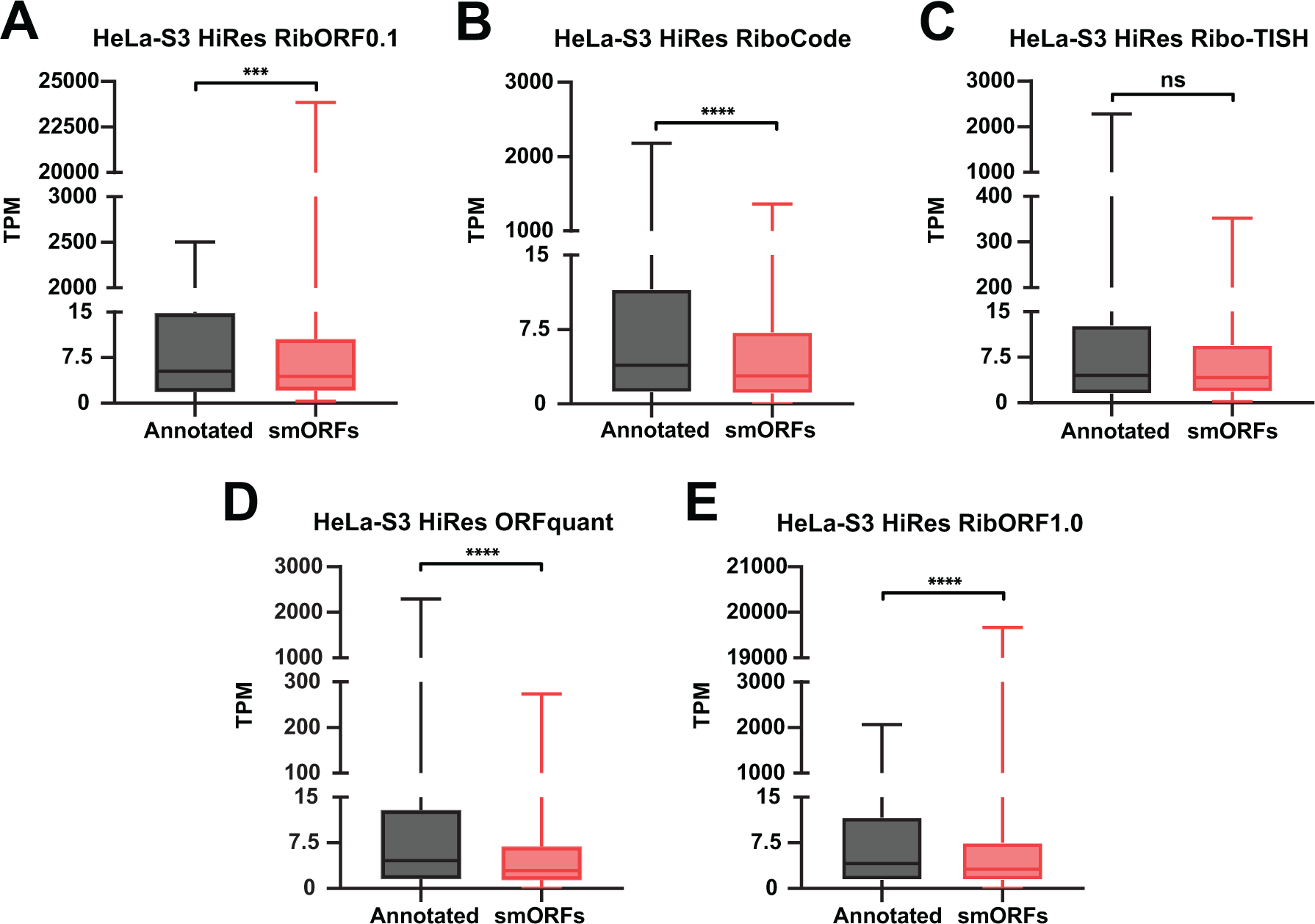
Comparison of Ribo-seq coverage in annotated genes and smORFs called translated in HeLa-S3 high-resolution dataset. (A-E) Quantification of Ribo-seq read coverage for both annotated gene ORFs and smORFs called translated by each tool in the HeLaS3 high-resolution dataset: RibORFv0.1 **(A)**, RiboCode **(B)**, Ribo-TISH **(C)**, ORFquant **(D)**, and RibORFv1.0 **(E)**. Coverage is calculated as transcripts per million (TPM) and are shown in Box-and-whisker plots. The box is bounded by the first and third quartiles, centerline shows the median, and whiskers represent the min and max values. Two-tailed Mann Whitney test was used to determine significant differences in coverage between annotated ORFs and smORFs (ns – not significant; ***, p < 0.001; ****, p < 0.0001).

**Supplementary Figure 10.**
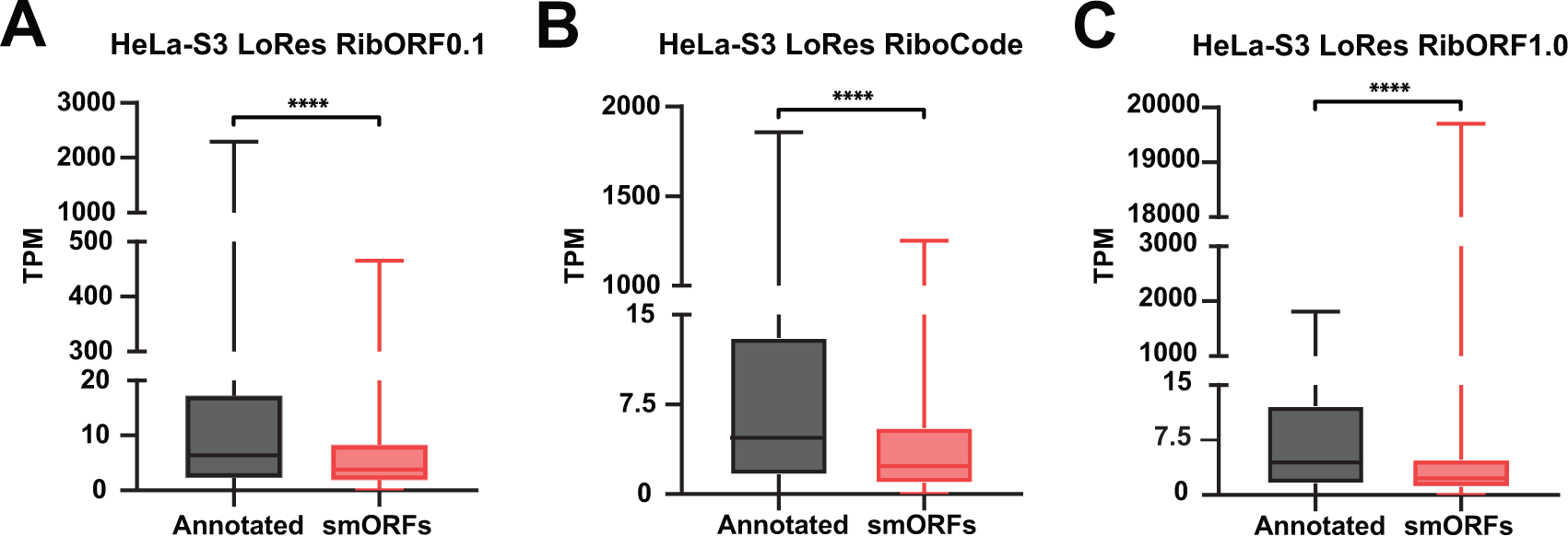
Comparison of Ribo-seq coverage in annotated genes and smORFs called translated in HeLa-S3 low-resolution dataset. (A-C) Quantification of Ribo-seq read coverage for both annotated gene ORFs and smORFs called translated by each tool in the HeLaS3 high-resolution dataset: RibORFv0.1 **(A)**, RiboCode **(B)**, and RibORFv1.0 **(C)**. Coverage is calculated as transcripts per million (TPM) and are shown in Box-and-whisker plots. The box is bounded by the first and third quartiles, centerline shows the median, and whiskers represent the min and max values. Two-tailed Mann Whitney test was used to determine significant differences in coverage between annotated ORFs and smORFs (****, p < 0.0001).

**Supplementary Figure 11.**
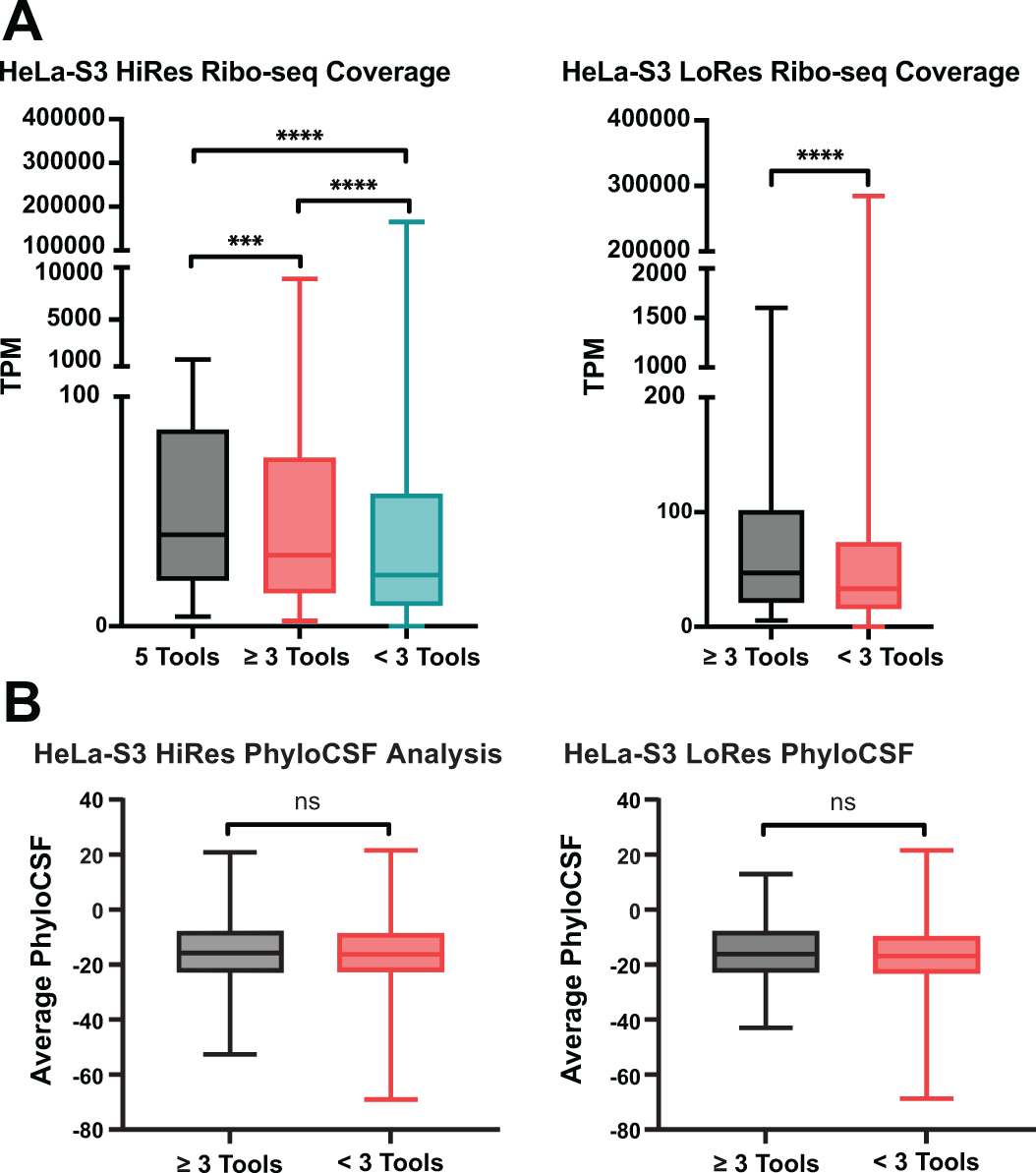
smORFs with higher Ribo-seq coverage are more likely to be predicted by multiple tools (A) Ribo-seq read coverage for smORFs predicted in high- and low-resolution HeLa-S3 datasets were categorized as identified in all five tools, in greater than three tools, and less than three tools. Coverage is calculated as transcripts per million (TPM) and are shown in Box-and-whisker plots. The box is bounded by the first and third quartiles, centerline shows the median, and whiskers represent the min and max values. For the low-resolution dataset analyses, no smORFs were found in common across all five tools. Two-tailed Mann Whitney test was used to determine significant differences in the Ribo-seq coverage (ns – not significant; ***, p < 0.001; ****, p < 0.0001). (**B)** Average PhyloCSF scores of smORFs predicted in high- and low-resolution HeLa-S3 datasets categorized by smORFs found in greater than three tools or less than three tools.

